# Exposure to epoxiconazole induces hyperlocomotion in adult zebrafish

**DOI:** 10.1101/2024.01.25.577223

**Authors:** Carlos G. Reis, Rafael Chitolina, Leonardo M. Bastos, Stefani M. Portela, Thailana Stahlhofer-Buss, Angelo Piato

## Abstract

Global agricultural production is sustained by an elevated use of pesticides. Their application in crops results in environmental contamination, which is demonstrated by widespread detections of these chemicals in aquatic ecosystems. This presence poses a risk to non-target organisms and ecological balance since the effects of potential exposure are misunderstood. Epoxiconazole (EPX) is a widely employed triazole fungicide, which has been frequently reported as a contaminant in superficial waters. However, the behavioral effects of EPX exposure in non-target organisms such as fish remain unexplored. We aimed to investigate the effects of EPX exposure on behavioral and biochemical outcomes, using zebrafish as a model organism. For this purpose, a static system was set with concentrations (24, 144, 240 μg/L) based on the fungicide environmental detection. The novel tank test (NTT) was performed after 96 h of exposure. The social preference test (SPT) was executed after 120 h, as well as the biochemical assays. In the NTT, the animals increased distance traveled, crossings, entries in the top area, and mean speed, indicating hyperlocomotion. No significant effects were observed in the SPT and the biochemical analyses. We suggest that the increased locomotion is related to the stimulant properties of EPX. Our results contribute to an in-depth understanding of this fungicide effects in zebrafish, which is essential for environmental contamination monitoring and management.

## Introduction

Constantly expanding, agriculture is one of the human activities that occupy the most territory on Earth (Zabel et al. 2019; Potapov et al. 2022). In 2021, the estimated cropland area accounted for 12.14% of the total terrestrial territory (FAOSTAT 2023), resulting in a primary commodities production reaching 9.5 billion tons (FAO 2022). This substantial output has various purposes, prominently including the nourishment of a global population exceeding 8 billion people (UN DESA 2022).

Along with the extensive occupied area, the widespread use of pesticides supports this remarkable production (Sharma et al. 2019). The misuse of these compounds poses an environmental risk, as their residues contaminate the ecosystems, leading to the impairment of non-target organisms (Dhuldhaj et al. 2023). The impact of these substances on the organism relies on its concentration, kinetics, mechanism of action, and the detoxification ability of the species (Escher et al. 2011). Simultaneously to the direct harm potentially induced by these compounds, the organisms can be affected through perturbations in the food chain, triggering ecological imbalance (Sarker et al. 2023). Contamination of superficial waters with pesticides constitutes a threat to aquatic species, such as fishes, since exposure to these chemicals affects their natural physiology (Subaramaniyam et al. 2023; Impellitteri et al. 2023). The pathophysiological effects of pesticide exposure include metabolic, reproductive, and neurobehavioral alterations (Bojarski and Witeska 2020; Cui et al. 2023).

Azoles are a widely used group of pesticides employed as fungicides (Jørgensen and Heick 2021). They are chemically characterized as compounds containing at least one nitrogen atom in a heterocyclic five-membered ring (Teixeira et al. 2022). When the ring contains three nitrogen atoms, these substances are specifically identified as triazoles (Matin et al. 2022). As a common mechanism of action, they inhibit the fungal enzyme lanosterol 14-alpha demethylase (CYP51), blocking the biosynthesis of steroids, mainly ergosterol (Huang et al. 2022). In non-target organisms, triazoles act as endocrine disruptors, interfering with the cytochrome P450 (CYP450) enzymes (Taxvig et al. 2008; Malcolm et al. 2009). Within this fungicide group, epoxiconazole (EPX) (C_17_H_13_ClFN_3_O) is commonly applied to control fungal diseases in both crops and seeds (Chambers et al. 2014; Wang et al. 2022). Considered stable in water, and exhibiting persistence under certain conditions (Lewis et al. 2016), likewise other pesticides EPX undergoes dissemination processes such as preferential flow and surface runoff, ultimately reaching aquatic ecosystems (de Castro Lima et al. 2020). The presence of EPX in surface waters is demonstrated by multiple detections across different locations worldwide, which threaten the species and the balance of the ecosystems (Guarda et al. 2020; Vera-Candioti et al. 2021; Zhang et al. 2023).

Studies have experimentally demonstrated the effects of exposure to EPX in non-target organisms. In an investigation involving the crustacean *Daphnia magna*, a concentration of 25 μg/L led to an increase in the protein content of adults within 1 to 3 days. Furthermore, there was an increase in the cumulative number of offspring when the exposure duration exceeded 31 days (Gottardi et al. 2017). In rats, oral exposure at doses of 8, 24, 40, and 56 mg/kg bw/day for 28 days caused neurotoxicity, showing alterations in outcomes linked to oxidative stress, DNA damage, and histology of the brain (Hamdi et al. 2022b). Similarly, the exposure to EPX (1.75 μg/kg bw/day) in female pregnant mice (FO) induced transgenerational effects persistent until the F3 generation, represented by phenotypic, histological, and transcriptomic alterations in the liver (Le Corre et al. 2022). The toxicity of this fungicide (100 and 1000 μg/L) was also verified in adult zebrafish, concluding altered energy, lipid, and amino acid metabolism, as well as histopathological changes in the liver (Jia et al. 2019). Additionally, exposure of embryonic zebrafish in concentrations of 175, 350, and 700 μg/L of EPX for 7 days induced an increase in the triglyceride and a decrease in glucose levels (Weng et al. 2021).

However, despite the existing evidence, some effects of EPX exposure in zebrafish were not fully elucidated. Specifically, studies describing possible behavioral alterations at environmental concentrations in adult zebrafish were never performed. Given the confirmed presence of EPX in the environment, it is essential to fill this gap to better clarify its impact on ecosystems. Therefore, this study aimed to investigate the effects of EPX exposure in adult zebrafish concerning behavior and biochemical outcomes.

## Methods

### Animals

Experiments were performed using 192 male and female (50:50 ratio) short-fin heterogeneous wild-type adult zebrafish (*Danio rerio*), approximately 6 months old, and weighing 300 to 500 mg. Animals were obtained from a local commercial supplier (Flower Pet, RS, Brazil). Fish were housed in 16-L (40 × 20 × 24 cm) unenriched glass tanks at a maximum density of 5 animals per liter. The water was dechlorinated using sodium thiosulfate pentahydrate 10% w/v (Na_2_S_2_O_3_ · 5H_2_O) (Neon, SP, Brazil). Tank water kept the controlled conditions required for the species (26 ± 2 ºC; pH 7.0 ± 0.3; dissolved oxygen at 7.0 ± 0.4 mg/L; total ammonia at <0.01 mg/L; total hardness at 5.8 mg/L; alkalinity at 22 mg/L CaCO_3_; and conductivity of 1,500–1,600 μS/cm), and was constantly filtered by mechanical, biological, and chemical filtration systems. Food was provided twice a day with commercial flaked food (Poytara, SP, Brazil) and *Artemia salina*.

Following AVMA Guidelines for the Euthanasia of Animals, the animals were euthanized by rapid chilling (2 −4 °C) for at least 10 minutes after cessation of opercular movement and decapitation to ensure death (AVMA 2020).

### Chemicals

Epoxiconazole (PESTANAL®, analytical standard, racemate) (CAS 133855-98-8) was acquired from Merck® (Darmstadt, HE, GER). Epoxiconazole was diluted in dimethyl sulfoxide (DMSO, CAS 67-68-5) (Sigma-Aldrich, St. Louis, MO, USA) (final concentration of 0.024% DMSO). Other reagents for biochemical analyses were obtained from Sigma-Aldrich®.

### Experimental procedures

The exposure was based on the guidelines of the Organisation for Economic Co-operation and Development (OECD) for acute fish toxicity tests (OECD 203) (OECD 2019). The concentrations were derived from a study that identified a maximum concentration of 144 μg/L of EPX in surface waters (Vera-Candioti et al. 2021). This concentration was used as the central value on the curve, with 24 μg/L and 240 μg/L representing the lower and upper extremes, respectively (24, 144, and 240 μg/L). Fish were exposed to epoxiconazole under static conditions in 4-L static tanks (17 × 17 × 17 cm) containing only an aeration system. Two tanks for each experimental group were used to eliminate potential tank effects. The resulting eight tanks were paired in the water bath apparatus. The compound was added once, and the animals were exposed for 96 h or 120 h depending on the test performed. During this period, the animals were monitored for injury or death and the feeding and temperature control were performed as usual. The allocation of the experimental groups was randomized and the researchers who directly handled the animals were blinded to the identification of the tanks. Researchers who did not handle the animals nor perform the analysis of the results performed the codification. The codes of experimental groups were revealed only during the statistical analysis. The sex of the animals was confirmed after euthanasia by dissecting the gonads. Figure 1 shows the experimental design.

**Figure 1.**
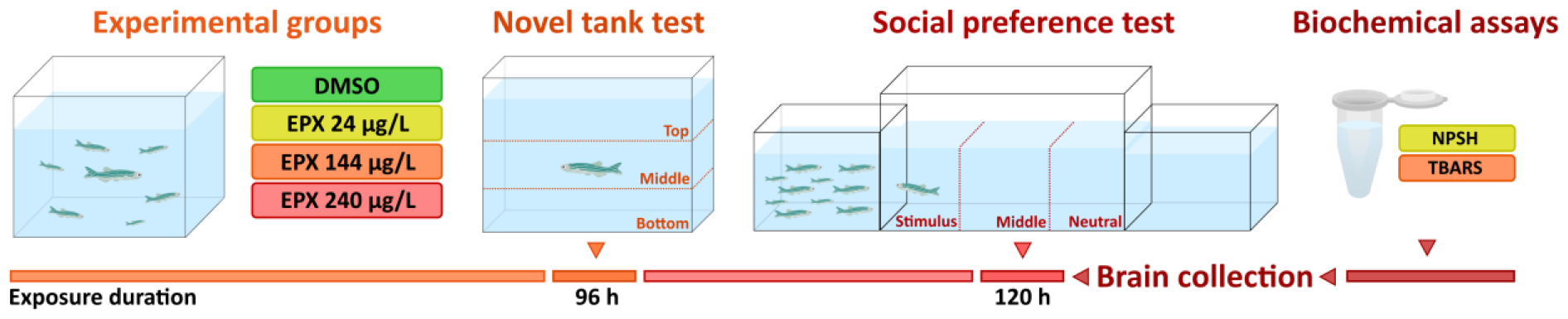
Experimental design. DMSO = dimethyl sulfoxide. EPX = epoxiconazole. NPSH = non-protein thiols. TBARS = Thiobarbituric acid reactive substances.

### Behavioral tests

#### Novel tank test (NTT)

After 96 h of exposure, animals were individually placed in the NTT apparatus (2.7-L tanks, 24 x 8 x 20 cm, 15 cm water level) and video recorded for 6 min from the front view. The software ANY-maze® (Stoelting Co., USA) was used to analyze the movement of the animals. The water of the test tank was renewed between each animal to avoid interferences. Locomotor and anxiety behaviors were measured by the total distance traveled (m), number of crossings (transitions among areas), time spent (s) and number of entries in the top area of the tank, mean speed (m/s), maximum speed (m/s), and immobility time (s) and episodes (Reis et al. 2020; Bertelli et al. 2021).

### Social preference test (SPT)

After 120 hours of exposure, animals were individually placed in a tank (30 x 10 x 15 cm, 10 cm water level) for 7 minutes. The tank was flanked by two identical tanks (15 x 10 x 13 cm, 10 cm water level), with one containing only water (neutral) and the other containing 10 unfamiliar zebrafish (social stimulus). The position of the social stimulus (right or left) was counterbalanced throughout the tests. The central tank was virtually divided into three vertical areas (stimulus, middle, and neutral). Videos were recorded from the front view. Animals were habituated to the apparatus for 2 min and then analyzed for 5 min. The software ANY-maze® (Stoelting Co., USA) was used to analyze the movement of the animals. The following outcomes were quantified: total distance traveled (m), number of crossings (transitions among areas), and time spent in the stimulus area (s) (as an index for social preference time). The water of the test tank was renewed between each animal to avoid interferences (Seibt et al. 2011; Benvenutti et al. 2020; Sachett et al. 2022). After the SPT, animals were euthanized for the biochemical assays described below.

### Biochemical assays

For each independent sample, four brains were collected and pooled (n = 6). The samples were homogenized in 600 µL of phosphate-buffered saline pH 7.4 (PBS, Sigma-Aldrich®) and centrifuged at 10,000 g, 4 ºC for 10 min in a refrigerated centrifuge; the supernatants were collected and kept in microtubes at −80 ºC until the execution of the assays. The detailed protocol for preparing brain tissue samples is available at protocols.io (Sachett et al. 2020c). The protein was quantified according to the Coomassie blue method using bovine serum albumin (Sigma-Aldrich, St. Louis, MO, USA) as a standard (Bradford 1976). The detailed protocol for protein quantification is available at protocols.io (Sachett et al. 2020a).

### Non-protein thiols (NPSH)

The content of NPSH in the samples was determined by mixing equal volumes of the brain tissue preparation (50 µg of proteins) and trichloroacetic acid (TCA, 6%), centrifuging the mix (10,000 g, 4 ºC for 5 min). Next, the supernatants were added to potassium phosphate buffer (TFK 1 M) and 5,5′-dithiobis-(2-nitrobenzoic acid) (DTNB 10 mM), and the absorbance was measured at 412 nm after incubation in the dark at room temperature for 1 h. The detailed protocol is available at protocols.io (Sachett et al. 2021).

### Thiobarbituric acid reactive substances (TBARS)

The levels of lipid peroxidation were assessed by TBARS assay. Samples (50 μg of proteins) were mixed with a solution of thiobarbituric acid (TBA 0.5%) and trichloroacetic acid (TCA 20%) (150 µL). The mixture was heated in a dry bath at 100°C for 30 min. The absorbance was determined at 532 nm in a microplate reader. Malondialdehyde (MDA 2 mM) was the standard. The detailed protocol is available at protocols.io (Sachett et al. 2020b).

### Statistical analysis

The sample size was calculated using G^*^Power 3.1.9.7 (Heinrich-Heine-Universität Düsseldorf, Düsseldorf, GER) for Windows according to the following criteria: one-way analysis of variance (ANOVA); effect size (0.4); alpha (0.05); power (0.9); number of groups (4). The total distance traveled was defined as the primary outcome. The total sample size was 96, resulting in n = 24.

It was used one-way analysis of variance (ANOVA) or the Kruskal-Wallis tests (Vrbin 2022). Outliers were defined by the ROUT statistical test using the distance traveled as the reference outcome. This resulted in one outlier being removed from the analyses in the NTT test, from the EPX 240 μg/L experimental group. The sex effect was tested in all comparisons demonstrating no effects. Therefore, the data were pooled.

Data are expressed as mean ± standard deviations of the mean (SD) or median ± interquartile range. A significance level of p<0.05 was considered.

Raw data are available at https://osf.io/a47hk/.

## Results

### Behavioral tests

#### Novel tank test

Figure 2 shows the results of the NTT. EPX 240 μg/L increased the distance traveled (F_3,91_ = 5.912, p = 0.0010, Fig. 2a), crossings (F_3,91_ = 4.350, p = 0.0065, Fig. 2b) and entries in the top area (F_3,91_ = 5.754, p = 0.0012, Fig. 2d), indicating hyperlocomotion. The outcomes of time in the top area (Fig. 2c), entries (Fig. 2e) and time in the bottom (Fig. 2f), immobility time (Fig. 2g), and immobility episodes (Fig. 2h) were not altered by any concentration.

**Figure 2.**
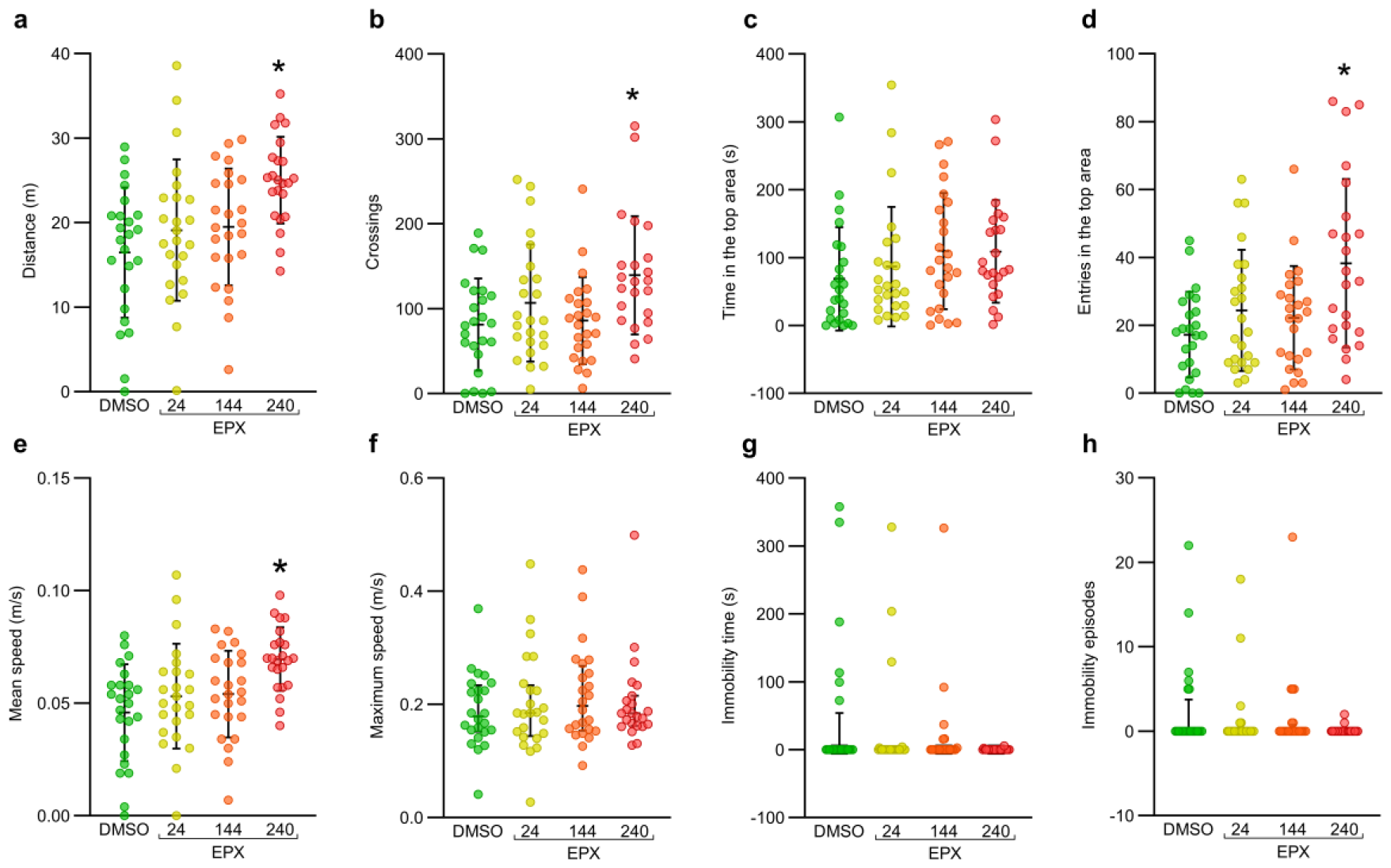
Effects of the exposure to EPX in the NTT. (a) distance, (b) crossings, (c) time in the top area, (d) entries in the top area, (e) mean speed, (f) maximum speed, (g) immobility time, and (h) immobility episodes. Data are expressed in mean ± SD and analyzed by one-way ANOVA followed by Tukey’s post hoc test (Fig. 2a-e) and as median ± interquartile range and analyzed by Kruskal–Wallis followed by Dunn’s test (Fig. 2f-h). Concentrations as µg/L. ^*^p<0.05 x DMSO. n = 24, except for EPX 240 µg/L n = 23. DMSO = dimethyl sulfoxide, EPX = epoxiconazole.

### Social preference test

Figure 3 shows the results of the SPT. At the tested concentrations, EPX did not induce alterations in any of the investigated outcomes.

**Figure 3.**
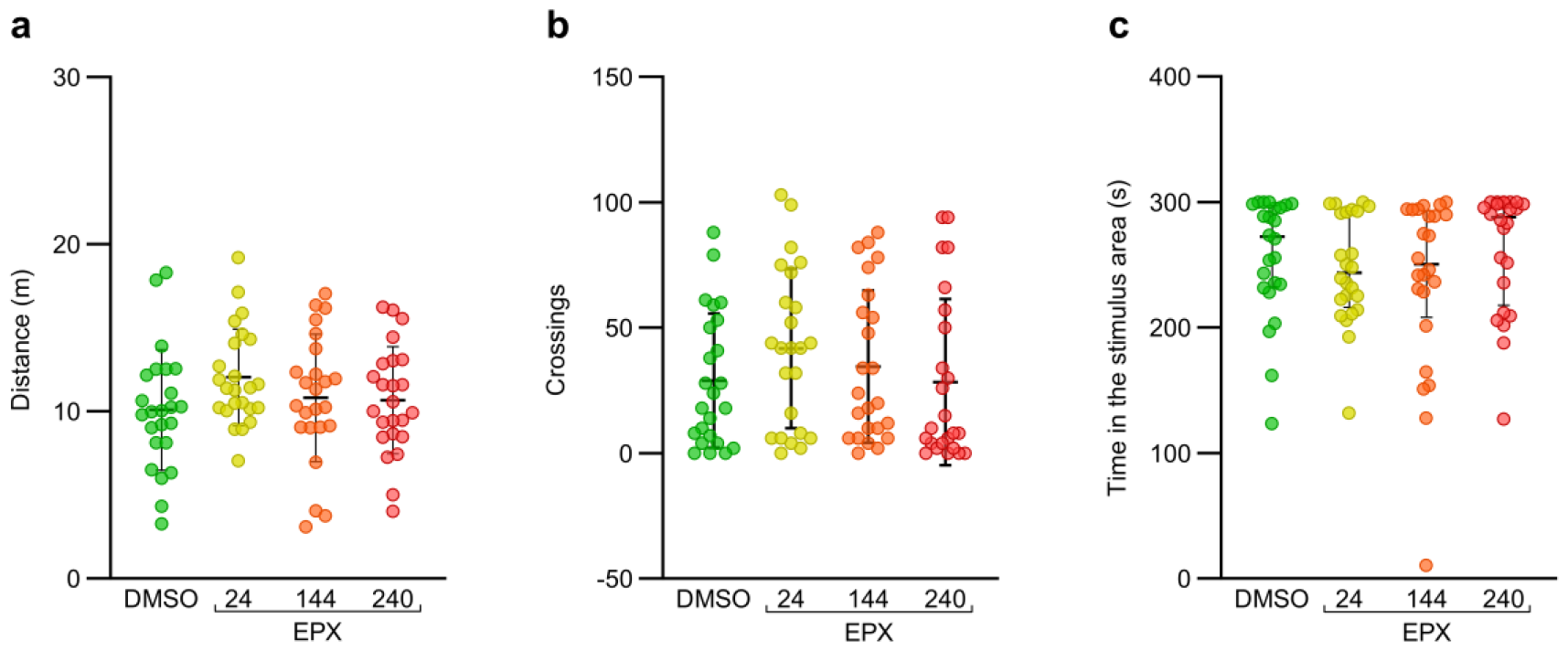
Effects of the exposure to EPX in the SPT. (a) distance, (b) crossings, (c) time in the stimulus area. Data are expressed in mean ± SD and analyzed by one-way ANOVA followed by Tukey’s post hoc test (Fig. 2a, b) and as median ± interquartile range and analyzed by Kruskal–Wallis followed by Dunn’s test (Fig. 2c). Concentrations as µg/L. n = 24. DMSO = dimethyl sulfoxide, EPX = epoxiconazole.

### Biochemical assays

Figure 4 shows the results of NPSH (Fig. 4a) and TBARS (Fig. 4b). EPX at the investigated concentrations did not alter either of these outcomes.

**Figure 4.**
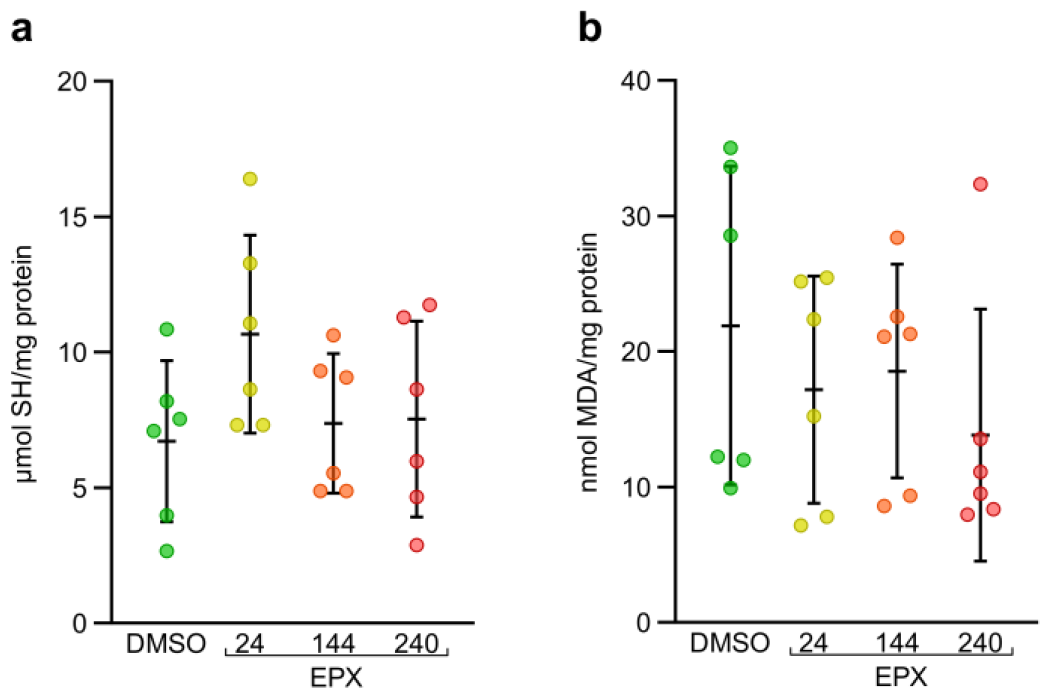
Effects of the exposure to EPX in NPSH and TBARS. (a) NPSH, (b) TBARS. Data are expressed in mean ± SD and analyzed by one-way ANOVA followed by Tukey’s post hoc test. Concentrations as µg/L. n = 6. DMSO = dimethyl sulfoxide, EPX = epoxiconazole, MDA = malondialdehyde, SH = thiols.

The results of statistical analyses for each outcome are detailed in Table 1.

**Table 1.** Results of statistical analyses for each investigated outcome. Significant effects (p<0.05) are given in bold font. DF = degrees of freedom. NPSH = non-protein thiols. TBARS = thiobarbituric acid reactive species.

| Test | Dependent variable | F-value or Kruskal-Wallis H | DF | p-value |
| --- | --- | --- | --- | --- |
| Novel tank | Distance traveled | 5.912 | 3,91 | **0.0010** |
|  | Crossings | 4.350 | 3,91 | **0.0065** |
|  | Time in the top area | 1.382 | 3,91 | 0.2533 |
|  | Entries in the top area | 5.754 | 3,91 | **0.0012** |
|  | Mean speed | 5.914 | 3,91 | **0.0010** |
|  | Maximum speed | 0.7980 | - | 0.8500 |
|  | Immobility time | 3.192 | - | 0.3630 |
|  | Immobility episodes | 3.112 | - | 0.3747 |
| Social preference | Distance traveled | 1.396 | 3,92 | 0.2490 |
|  | Crossings | 1.007 | 3,92 | 0.3935 |
|  | Time in the stimulus area | 3.994 | - | 0.3637 |
| NPSH | $\mu\text{mol SH/mg protein}$ | 1.787 | 3,20 | 0.1820 |
| TBARS | $\text{nmol MDA/mg protein}$ | 0.7521 | 3,20 | 0.5340 |

## Discussion

This study aimed to evaluate the effects of exposure to epoxiconazole at environmental concentrations in adult zebrafish. As the main findings, we can highlight that epoxiconazole increased the total distance traveled, crossings, entries in the top area, and mean speed in the NTT. Taken together, these alterations are evidence of hyperlocomotion. In contrast, no effects were observed in the SPT or the biochemical analyses.

There are relatively few studies in the literature concerning the behavioral effects of exposure to EPX. Even in widely used model organisms such as rodents, the investigation of behavioral outcomes is unusual. Other toxic effects are more often reported in these mammals, such as hepatotoxicity, reproductive and developmental toxicity, and cardiotoxicity (de Castro and Maia 2012; Hester et al. 2012; Schneider et al. 2013; Hamdi et al. 2022a). The same pattern in the literature is verified for non-zebrafish fish species, where few studies with these aquatic animals are available. Although there are some available studies of exposure to EPX in zebrafish, none of them explored its potential consequences regarding the behavior in different domains (Reis et al. 2023).

Distance traveled and mean speed are some of the endpoints used as indicators of locomotor behavior in the NTT in zebrafish (Kysil et al. 2017). In our study, EPX exposure increased the distance, crossings, entries in the top area, and mean speed, which suggests hyperlocomotion. Our findings contradict what was reported by a recent metanalysis (Reis et al. 2023). The study showed that fungicides exposure overall and triazoles specifically decrease the distance traveled both in larvae and adult zebrafish. However, this systematic review did not include any study with EPX. Locomotor behavior is fundamental for vertebrates to survive and maintain basic functions, such as foraging, feeding, avoiding a threat, and breeding (Berg et al. 2018). It is characterized as a complex behavior, which depends on the integration of the brain and spinal cord (Grillner and El Manira 2020). In this network, the neurotransmitter dopamine has a prominent role in the activation and modulation of the locomotor system across animal species (Sharples et al. 2014). Likewise, the dopaminergic system is directly linked to swimming in zebrafish (Matsui 2017). This functional relation is proven by pharmacological studies, in which the exposure to D_1_ and D_2_ selective agonists caused locomotor hyperactivity in zebrafish larvae, whereas selective antagonists of the same receptors induced hypoactivity (Irons et al. 2013). Additionally, the exposure to the triazole fungicide triadimefon (20 mg/L) induced hyperlocomotion in adult zebrafish due to its property of inhibition of dopamine reuptake (Paredes-Zúñiga et al. 2019). In another study, this compound (5 mg/L and 15 mg/L) increased the circling behavior (Paredes-Zúñiga et al. 2021). Besides the typical mechanisms of toxicity shared by all triazoles, like the interference in CYP450 enzymes, triadimefon also causes the accumulation of synaptic dopamine by binding in the dopamine transporter (DAT) (Moser 2014). Even though in a speculative view, we suggest that the observed hyperlocomotion effects of EPX 240 μg/L in this study are due to an increase in dopamine levels. Considering that these compounds, EPX, and triadimefon, belong to the same chemical group, it is feasible that EPX could present still unknown similar properties. The same argument can be used to clarify our divergence from the above-cited metanalysis (Reis et al. 2023).

Regarding the SPT, no effects of EPX exposure were observed. In this test, the social preference can be assessed by the attempt of the animal to stay near to his conspecifics, which is measured by the time spent in the stimulus area (Benvenutti et al. 2020). Notably, there is a gap in the literature about the potential effects of EPX exposure on social behavior, in both rodents and zebrafish. Social behavior holds significant importance for the survival of zebrafish, involved in foraging, shoaling, schooling, and antipredator behavior (Müller et al. 2017). The innate and robust nature of the social preference suggests that considerable interventions may be required to disrupt it (Pagnussat et al. 2013; Ogi et al. 2021). The duration of exposure and concentrations of EPX used in this study, appear to be insufficient to elicit abnormal behavior in this test.

Concerning the biochemical analyses, no effects were observed in NPSH and TBARS levels after 120 h of exposure. Other studies showed that the exposure to EPX at daily doses of 8, 24, 40, and 56 mg/kg for 28 days increased the level of MDA relative to the TBARS assay in the kidney, liver, and brain of rats, indicating lipid peroxidation (Hamdi et al. 2019, 2022b). Moreover, the embryos of rare minnow (*Gobiocypris rarus*) exposed to EPX for 72 h showed an increase (2 mg/L) and decrease (5 mg/L) in the activity of superoxide dismutase (SOD), while both concentrations decreased glutathione S-transferase (GST), adenosine triphosphatase (ATPase), and acetylcholinesterase (AChE) activities (Zhu et al. 2014). Since the duration of exposure and concentrations used in this study were not able to cause detectable oxidative damage, we did not quantify antioxidant enzymes. Even if we detected alterations in enzymatic activity, it would probably be due to successful physiological processes that are fulfilling the aim of homeostasis preservation. Thus, it is likely that the hyperlocomotion is not linked to oxidative damage.

As limitations of this study, we firstly emphasize the use of analytical-grade EPX instead of a commercial formulation. This decision was deliberate to investigate the toxicological effects of the isolated fungicide, which would be impossible with commercial products since they contain other ingredients. Frequently, these other substances are undisclosed by the manufacturers. We acknowledge that this is a less realistic approach, once the pesticides used in agriculture come from a commercial source. Furthermore, pesticides are present in the environment as a mixture with other contaminants, which makes the analysis of the combined and synergistic effects more coherent. A comparative study with both formulations, analytical-grade, and commercial, is needed to better characterize the effects of EPX in the environment, as well as substance combinations. In the environment, the duration of exposure is extremely dynamic and potentially long-lasting, and the delimited 96 and 120-hour periods used in this study did not reflect these conditions. Finally, we did not perform experiments with exposure in embryonic and larval zebrafish, so we cannot suggest if there are different effects according to the developmental stage. It is reasonable that a potential exposure starts earlier in the life cycle of the animal.

In this study, we described hyperlocomotor effects caused by EPX exposure in zebrafish, without any observable alterations in social and biochemical parameters. Locomotor alterations pose a risk to the survival of the species, considering that it is a core behavior to comply with basic demands. Once an organism is affected, it can compromise the balance of an entire ecosystem. Due to the increasing evidence of the presence of emergent contaminants in the aquatic environment, our findings contribute to a comprehensive view of the consequences of this contamination. This knowledge can be useful to elaborate both monitoring and management strategies for environmental pollution.

## Statements and Declarations

### Data availability

The datasets used and analyzed during the current study are available at Open Science Framework (https://osf.io/a47hk/).

### Author contributions

All authors had full access to the data in the study and took responsibility for its integrity and the accuracy of the data analysis. Conceptualization: Carlos G. Reis and Angelo Piato; Formal analysis: Carlos G. Reis; Funding acquisition: Angelo Piato; Investigation: Carlos G. Reis, Rafael Chitolina, Leonardo M. Bastos, Stéfani M. Portela, Thailana Stahlhofer-Buss; Methodology: Carlos G. Reis and Angelo Piato; Project administration: Carlos G. Reis; Resources: Angelo Piato; Supervision: Angelo Piato; Visualization: Carlos G. Reis, Leonardo M. Bastos; Writing - original draft: Carlos G. Reis; Writing - review and editing: Rafael Chitolina, Leonardo M. Bastos, Stéfani M. Portela, Thailana Stahlhofer-Buss, Angelo Piato. All authors read and approved the final manuscript.

### Ethics approval

All procedures were approved by the Universidade Federal do Rio Grande do Sul ethical committee (#37711/2019) and were performed following relevant guidelines on the care and use of laboratory animals and following Brazilian legislation regarding animal research.

